# A new Image Segmentation Algorithm with Applications in Confocal Microscopy Analysis

**DOI:** 10.1101/524389

**Authors:** M. Sánchez-Aragón, F. Casares

## Abstract

Gene regulatory networks (GRNs) represent the molecular interactions that govern the behavior of cells in tissues during development. The building and analysis of GRNs require quantitative information on gene expression from tissues. Laser Scanning Confocal Microscopy (LSCM) is commonly used to obtain such information, where immunofluorescence signal can be used as a correlate of gene expression or protein levels. However, a critical step for the extraction of this information is the segmentation of LSCM digital images. Popular segmentation algorithms are frequently based on watershed methods. Here we present an algorithm for the 3D segmentation of *nuclei* from LSCM (x,y,z) image stacks based on regional merging and graph contractions. This algorithm outperforms watershed methods, especially when the density of images along the z-axis is low and there is a high nuclear signal crowding. In addition, it reduces the parameterization since no filter is needed in order to prevent signal noise side effects (e.g. oversegmentation). Based on this algorithm, we developed an application (iFLIC, immunoFLuorescence Imaging Cytometry tool) for the Java Virtual Machine (JVM). The application supports basic operations for reading, writing and filtering 8-bit depth multicolor TIFF image formats, including indexed file directories (IFD), which are provided by the Java Advanced Imaging (JAI) library. It also provides with basic 3D-rendering and ROI specification that make extensive use of the Java3D library. iFLIC is also a plugin based application powered by the Java Plugin Platform (JPF), so each specific operation is declared as a unique command associated to one plugin and linked to a common interface. Results from segmentation can be exported both as TIFF images and a descriptive file format (iFLIC format)

## 1 introduction

The functional analysis of DNA elements and the study of the role of specific transcription factors (TF) in the control of gene expression during *metazoa* ontogenesis has led to description of gene regulatory networks (GNR) which is allowing the design of quantitative models for their study. The analysis of the critical properties from these models, such as ergodicity, sensitivity to initial conditions, noise robustness or oscillatory and chaotic behaviors, are of major importance for their validation, for which a quantitative description of the evolution of gene expression in space and time is necessary at a cell resolution level. Laser scanning confocal microscopy (LSCM) has proved to cope well with the need of this detailed description in developmental tissues, due to its sensitivity, resolution and wide distribution in imaging facilities. However the extraction of meaninful quantitative information requires segmentation procedures for automatic high-throughput retrieval of image features. In the study of GRNs, since these are operated by nuclear transcription factors, the focus of these procedures is the detection and extraction of *nuclei*.

The quantification of gene expression (using as proxy immunofluorescence intensity) using LSCM data and image segmentation algorithms and their application in GRN reconstruction has been pioneered by studies of the early patterning of *Drosophila* embryos. This has been so because *Drosophila* blastoderm *nuclei* are superficially located in the embryo, uniformly spread out and undergo synchronic cell divisions in a stereotypic manner [1]. The data obtained from these embryos has allowed the maturation of dynamical models that contributed to the characterization of the quantitative effects on regulation for classical early genes such as *Krüppel* (*Kr*) or *even-skipped* (*eve*) [2, 3].

Beside blastoderm, *Drosophila* also offers the opportunity for studying developmental processes in other tissues, among which remarkable examples are the *imaginal discs*. the *Drosophila* imaginal discs are the larval precursors of adult structures, such as the wings or eyes [4]. They are also very powerful models to understand developmental processes, and a number of efforts have been directed towards the building of GRNs explaining their patterning and growth [5]. However, unlike blastoderms, imaginal discs are more challenging for image analysis: although they are formed by a single epithelium, this epithelium is pseudostratified (i.e. the *nuclei* are not located on a single plane), presents folds, and stained *nuclei* are small and densely packed (figure 1), which makes nuclear segmentation much more difficult. In particular, the motivation of our research has been the quantitative analysis of gene expression during the development of the eye disc. The difficulty in obtaining detailed information from this tissue has precluded the quantitative modeling of the *Drosophila* eye GRN, despite the fact that a GRN has been already been described using genetic and molecular biology techniques [6].

**Figure 1:**
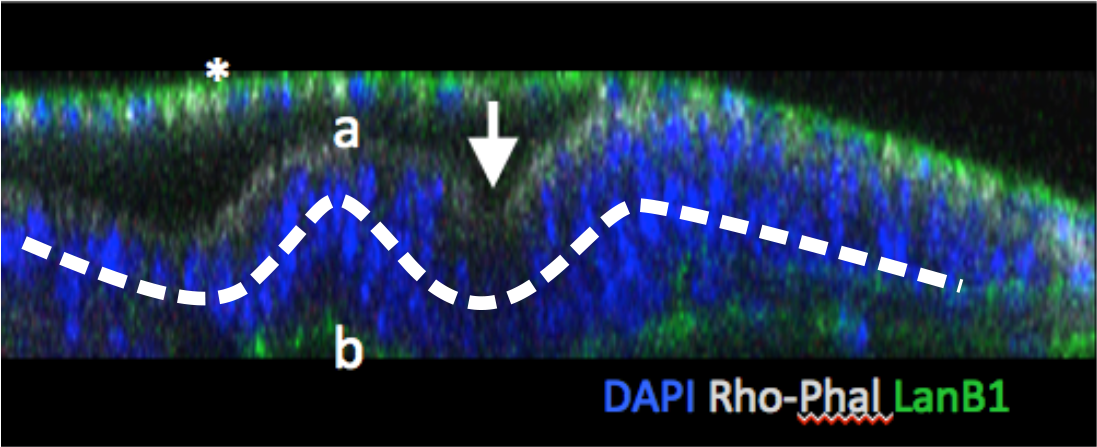
x,z confocal section of an eye disc stained for nuclei (DAPI; blue), apical actin (Rhodamine-phalloidin, white), basal membrane (lamininB1:GFP, green). “a” and “b” indicate the apical and basal sides of the epithelium. The arrow marks the morphogenetic furrow. The asterisk (“*”) marks the the thin periodical epithelium that covers the eye epithelium.

Most common segmentation methods are variations of the *watershed* algorithm, for which the reader may consult the excellent work of Cousty *et al.* [7, 8]. Watersheds have become popular for their easy implementation, low computational cost and three dimensional extensibility. Watersheds follow the intuitive idea that overlaps in crowded *nuclei* images could be solved by finding those regions that separate two grayscale minima, as in Cousty’s “drop-of-water principle”. In practice watersheds appear very sensitive to noise and irregular signals (e.g. DAPI signal often reveals nuclear structures that impairs the segmentation performance) leading to oversegmentation. The use of filters and thresholding subroutines has been aimed at improving watershed methods performance [9, 10], for which available implementations can be found [11, 12, 13]. However the effectiveness of the pipeline is still not satisfactory in all cases. Azuma & Onami tested the performance of different variations for the watersheds algorithms during the development of *Caenorhabditis elegans* in the prescence and absence of a differential of gaussians filter (DoG) [14]. Their results indicate that error rates were acceptable up to a 5 per cent of significance until the stage of 350 cells. Beyond this point the error rate increased according to tissue complexity, insofar as similarities with manual segmentation were significantly divergent.

Other authors have implicitly recognized the problem that LCSM entails for the assignation of segmented regions accross different confocal planes, for which methods have been proposed [15, 9]. We approached the problem of the segmentation of LSCM images from *nuclei*-stained tissues of high complexity taking into account limitations in both LSCM and in watershed-based methods. On one hand we think LSCM should be regarded as a technique with a poor z-resolution for three dimensional reconstruction; for instance the relative resolution in the z direction is frequently so low relative to the other ortogonal directions that confocal planes should be taken as independent items with a sort of topological relation. This fact changes, not only how well we look at a three dimensional object (i.e. a problem of raw spatial resolution), but more importantly how much information is available to infer the connection between the visible pieces of each object, i.e. their intersection with confocal planes, something we may call *topological resolution*. On the other hand we think that, due to signal irregularities occurring naturally on biological samples (e.g. euchromatin features), boundary detecting approaches do not necessarilly lead to an accurate segmentation of *nuclei*, not even when the image is binarized since the features of the boundaries themselves are the ones that determine the output, requiring filters to be applied previously, and so increasing user parametrization a reducing reproducibility.

In this work we propose a new method for three dimensional segmentation of LCSM derived images based in a different concept: instead of focusing on boundary motivated procedures, we tried to describe the objects as a combination of connected subspaces, assuming that we only know with precision some parts of them: their intersection with confocal planes. For that we took advantage of topological definitions in the context of metric spaces and propose a regional merging procedure which results naturally in a reproducible segmentation that is in agreement with manually annotated images. The only assumed step previous to segmentation is the binarization of the image stack using an appropiate method such as an otsu [16], which may also be enhanced sometimes by performing histogram equalization [17].

## 2 methods

### 2.1 definition of the problem

Let *S* be some compact subspace, not composed of disjoint sets and contained in *T* ⊆ ℝ^*n*^, and consider the set *ζ* = {*B*_*i*_} = {*B*(*r*_*i*_, **p**_*i*_) ⊆ *S*}, i.e. the set of all closed *n*-balls fully contained in *S*, for some radius *r*_*i*_ ∈ ℝ and some point **p**_*i*_ ∈ *S*∀*i*, *n* ∈ ℕ. The ball 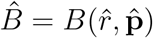 such that 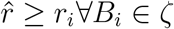 is the *maximum ball* contained in *S*.

Let ζ′ ⊂ ζ be a partition obtained by including recursively the maximum ball from each subspace 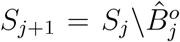 into one set, where 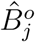 is the relative open ball of 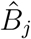, with *j* = 0 ⇒ *S*_*j*_ = *S*∀*j* ∈ {0, ℕ}. It can be seen that:

1. the union of ζ′ equals *S*
2. for any two 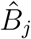, 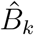 contained in ζ′, 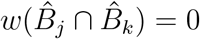, being *w* a function retrieving the extension or *weight* of a point set.

A key observation regarding the partition ζ′ is that for any pair of balls their intersection may not be empty despite that their volume may be zero (property two). In fact, for two typical balls in an euclidean space, a non-empty intersection whose volume is zero is some point in *S*. Property two allows us to represent the subspace *S* as an undirected graph *G*(*V, E*), for some set of vertexes *V* (equals to the elements of ζ′) and set of edges *E*, such that *E* includes all those pairs from ζ′ whose intersection is non-empty. We also impose *E* to be *weighted* in the sense that there exists an arbitrary map *ϕ* : *E* → ℝ. A minimum spanning tree (MST) from *G* under *ϕ* and their possible contractions are of major importance for the algorithm we are proposing later. In general *ϕ* may take any form depending on the properties of the vertex balls that belong to each edge, e.g. the values of *w*^1^. Regarding the properties of *G* we use two concepts in the following section, *degree reducing contraction* and *limit minor*, that we describe briefly below.

#### Definition 1 (degree reducing contraction)

*Consider G*′(*V, E*′ ⊆ *E*) *the minimum spanning tree of G under the map ϕ. Also define the degree δ of any vertex u*_*j*_ ∈ *V the sum of the weights of its nearest first order neighbours N*_*j*_, *and c*(*e*_*jk*_) *the resulting vertex from a contraction of an edge e*_*jk*_ = {*u*_*j*_, *u*_*k*_} ∈ *E*′, *u*_*k*_ ∈ *N*_*j*_. *e*_*jk*_ *is said to be degree reducing if*

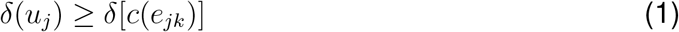

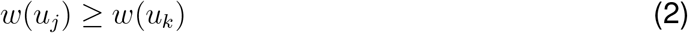

*The new vertex that arises from a contraction is considered to have a weight equal to the sum of the weights of the intervening sets*.

related to the degree reducing contraction we define the limit minor considering a recursive contraction of a weighted tree.

#### Definition 2 (limit minor)

*Consider some tree* 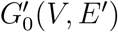 *with a set of vertexes V* = {*u*_*j*_} *that are labeled according to some function w and suppose there exists a set of edges* 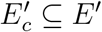 *(composed by the set of vertexes V*_*c*_ ⊆ *V) whose contractions are degree reducing. The contraction of some edge* 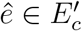 *that contains some* 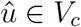 *as vertex such that* 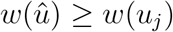 *for all u*_*j*_ ∈ *V*_*c*_ *results in the graph* 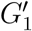. *By induction over* 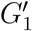 *a series of graphs* 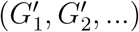 *is obtained, up to some graph* 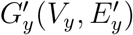 *for which the set of degree reducing edges is empty. We call this graph the limit minor of G′*.

We are interested in finding a sequence of degree reducing contractions of *G*′ that leads to its limit minor. From the list of elements of *V* we perform such contractions as long as the conditions 1 and 2 hold for at least one edge in *E*′. The process can be repeated until either condition is not satisfied for any set. The interest of the degree reducing edges come from the expectation that the union of the associated balls resembles the voxel aggegrates that typically define a *nucleus* in a set of confocal images, if an adecquate function *ϕ* is found, allowing a segmentation procedure by regional merging to be specified.

Finding a minor does not ensure that all initial regions are merged. In general just the subset of largest extension bear some interest. A classification method for the final sets and a function *ϕ* are proposed in the next section.

### 2.2 practical approach and algorithm outline

In practice the ideas explained previously cannot be used right away in order to segment *nuclei* coming from LSCM-derived images due to the fact that these images only account for some parts of the true objects. In other words, LSCM only offers an incomplete definition of the *nuclei*, i.e. their intersection with confocal planes. This creates an important uncertainty regarding the topological connectivity of all balls and hence how *G* is defined. We explored the possibility of formulating the problem in terms of balls intersections with the confocal planes, instead of using pure balls, and finding a function *ϕ*, among a finite set, such that it reproduces an adequate connectivity for the intended segmentation (figure 2).

**Figure 2:**
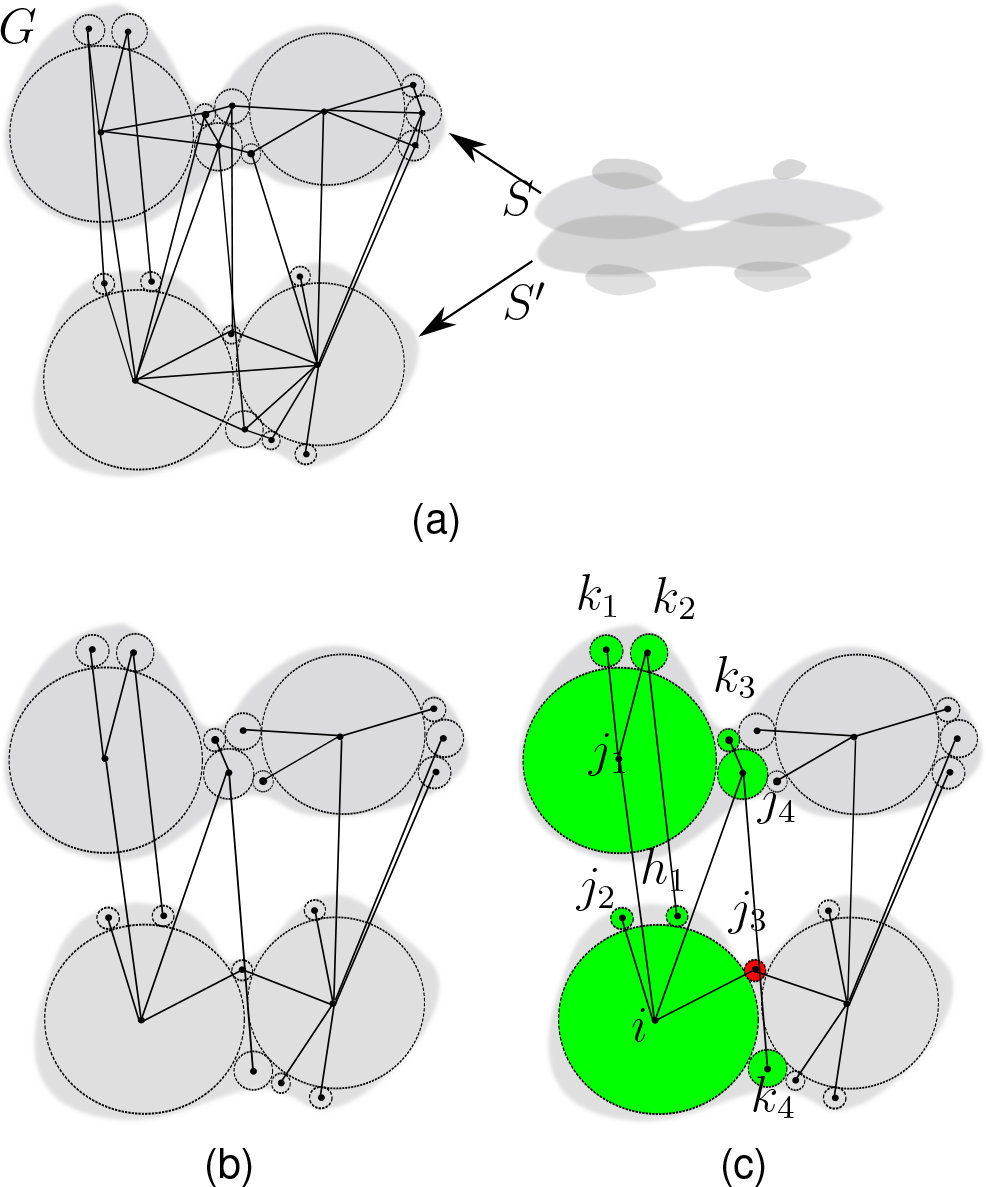
scheme of confocal plane connectivity and degree reducing paths. 2a initial graph. Maximum 3-balls intersections are shown for two adjacent confocal planes. Voxel adjacency defines a initial graph *G* for these intersections. 2b Resulting minimum spanning tree (MST) upon weighting the edges of *G* according to some function *ϕ*. 2c Reducing degree paths from the largest set (vertex) *i*, {< *i*, *j*_*x*_, *k*_*y*_, *h*_*z*_ >} either finish at some pendant set vertex(es) or at some set vertex(es) that does not satisfy the reducing degree condition (red, *j*_3_), which are generally considered as belonging to the *residual sets*. The other main vertexes associated to the paths define a partition of the graph *G* (green). The way we obtain such partitions is mainly imposed by *ϕ*

A well-defined domain, such as those that result from the limit minor *G*′, relies on the existence of degree reducing paths between adjacent balls in *G*′, but these paths were set up from a much wider set of edges of *G*. In order to obtain a result that it is more in agreement with visual experience we reasoned that if two ball intersections share a wide contacting area, e.g. z-projection overlap, their connection should be favoured since there is a higher probability of belonging to the same object. If the contacting area is reduced we favour a connection proportionally to the assymetry. This combination results in paths between wide contacting sets of points that satisfy some degree of assymetry, creating “bridges” at small sets of points, whose potential contact surface is limited. Following this idea we let ζ′ be composed of disjoint collections of voxels that represent a few set of plane intersections. *G*(*V, E*) is defined in terms of voxel adjacency: an edge is created between point sets *u*_*j*_ and *u*_*k*_ if some voxel from *u*_*j*_ has an immediate neighbour voxel from *u*_*k*_ located at any of the six orthogonal positions. *E* is weighted following the proposed map:

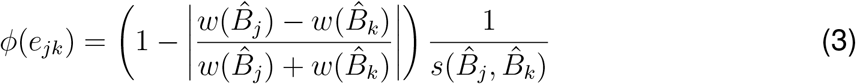

where *s* stands for the contact surface of 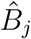 and 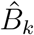, i.e., the number of voxels from either set of points that has at least one voxel from the other set of points as a neighbour.

The domains of degree reducing paths are defined in the same way as in the previous section from the MST of *G*. A limit minor of *G*′ is expected to render two groups of sets: those with large weight (i.e. extension), that account for most contractions of *G*′ and resemble quite well the shape of *nuclei*, i.e. *main sets*; connected by small sets of points, *residual sets*, that bridge up the main sets (figure 2c). We next use a hierarchical clustering procedure for grouping the final sets into these two classes. Hierarchical clustering is usually considered an optimization problem, where some cost function *f* is minimized in the process. This definition allowed the proliferation of methods that were adequate in suiting specific data sets depending of the function of choice [18]. Here we apply a variation of the Ward’s clustering where the cost function is the Fano factor [19]. Let ζ′ be split into two sets 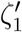 and 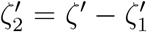. The cost of this split is determined by *f* and in our case it is expressed as:

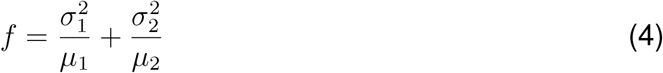

being 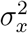 and *μ*_*x*_ the variance and mean of set weights in 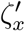.

This specifies the steps of an algorithm for the segmentation of each disjoint set of an image. Such an algorithm is sketched as follows:

1. obtain a binary image set, (e.g. otsu thresholding). This defines the compact subspace *S*
2. get the set of balls defined as a collection of plane intersections (ζ′).
3. build the initial graph *G*.
4. compute the minimum spanning tree *G*′ under the map 3.
5. find the limit minor of *G*′ by performing all their reducing degree contractions applying definition 1.
6. determine the residual sets by hierarchical clustering applying the cost function 4.
7. deliver the voxels from the residual vertexes among the main sets according to the proximity.

In our current implementation, maximum balls are obtained by checking signal matches of spheric patterns with decreasing radius in the whole stack of images. When a match is found, the inner voxels are removed from the image and included in a set. The process ensures that the retrieved regions are maximum balls of the complementary spaces (i.e., the set ζ′). For the construction of the graph *G* the boundary voxels for each set is determined, and a search into other set’s boundary performed. This step is usually time consuming since it tipically runs in a second-degree polynomial complexity 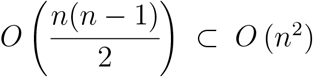, for any amount of balls *n*. We reduced the running time by limiting the number of possible neighbours proportionally to the number of its boundary voxels. Considering that a boundary voxel potentially bears 5 orthogonal neighbours^2^ the algorithm may run in expected time *O*_*E*_ (min{5〈*b*〉, *n* − 1}*n*) where 〈*b*〉 denotes the average size of the boundary. After *G* is specified, components are determined by Depth First Search (DFS) as described by Tarjan [20]. The MST from each graph component in *G* under the map 3 is calculated by an implementation of Prim’s algorithm [21], after which a limit minor is searched (algorithm 1). At this stage a priority queue is used in order to retrieve the maximum weighted ball of each iteration and its degree reducing edge. These edges undergo contraction and the spanning tree is updated with the new vertex. The operation is repeated until no further contraction is allowed for any ball.

After the DFS all graph components can be processed in parallel. Only at the end the final vertexes of the limit minor are collected from each component, undergoing hierarchical classification. The usage of Fano’s factor within the cost function ensures that the classification finds a point of minimum noise-to-signal ratio within the set of final vertexes, which we consider as a valid interpretation of the subdivision between the two kinds of sets by weight. Since the number of specified groups is prefixed to two, we may simply sort the sets by weight and find the minimum point of the cost function in one loop, by scanning from the smaller to the largest set.

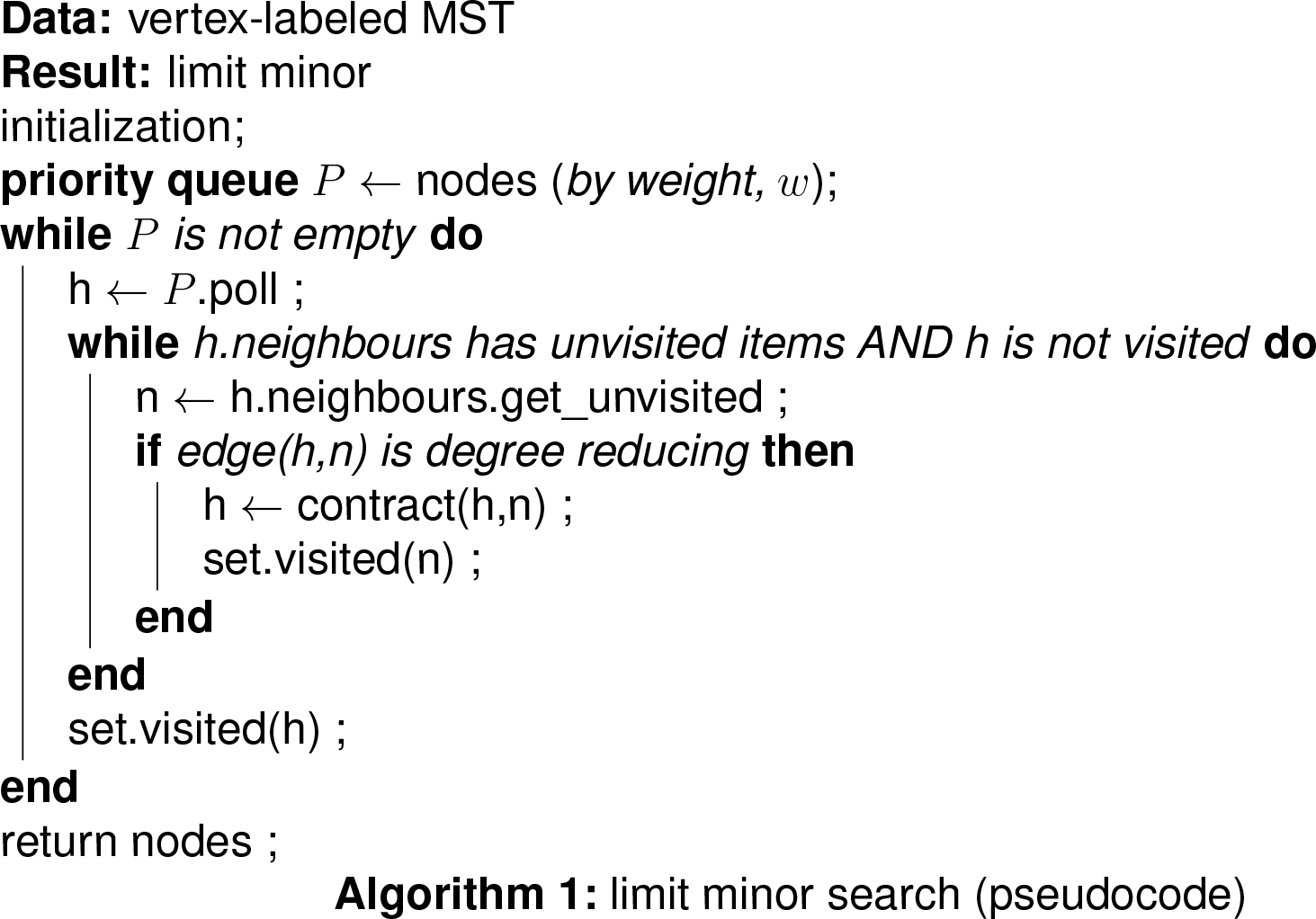

We developed an application (iFLIC, immunoFLuorescence Imaging Cytometry tool) for the Java Virtual Machine (JVM) that integrates the described algorithm. The application supports basic operations for reading, writing and filtering 8-bit depth multicolor TIFF image formats, including indexed file directories (IFD), which are provided by the Java Advanced Imaging (JAI) library. It also provides with basic 3D rendering and ROI specification that make extensive use of the Java3D library. iFLIC is a also a plugin-based application powered by the Java Plugin Platform (JPF), so that each specific operation is declared as a unique command associated to one plugin and linked to a common interface. Results from segmentation can be exported both as TIFF images and as a descriptive file format (iFLIC format). In order to provide functionality with data analysis applications, a package for R language is currently in development, from which some functions scripts may be sent upon request. The reader is encouraged to check the java code included in the appendix A for details.

## 3 results

In order to test the performance of the formalism described above, several stacks of images with manually drawn spots were generated in ImageJ and subjected to segmentation with iFLIC (figure 3). Spot voxels were forced to have the highest of two possible values, delimiting the signal level of the background. These spots were not demanded to have any concrete shape (e.g. spherical). We can observe that the thirteen voxel aggregates corresponding to these spots were correctly annotated under conditions of high signal crowding and overlap (figure 4). Sets of points composed by ball intersections with planes are connected by a criterium of voxel adjacency as described in the previous section. These voxel aggregates are generally dominated by large sets of points surrounded by small ones, imparting size gradients around and establishing paths of degree reducing-edges. After weighting the edges according to equation 3, the resulting MST is contracted up to its limit minor, leaving an alternating pattern of large sets connected by small sets. These sets are later sorted by hierarchical clustering into two groups using the cost function 4 (figure 5).

**Figure 3:**
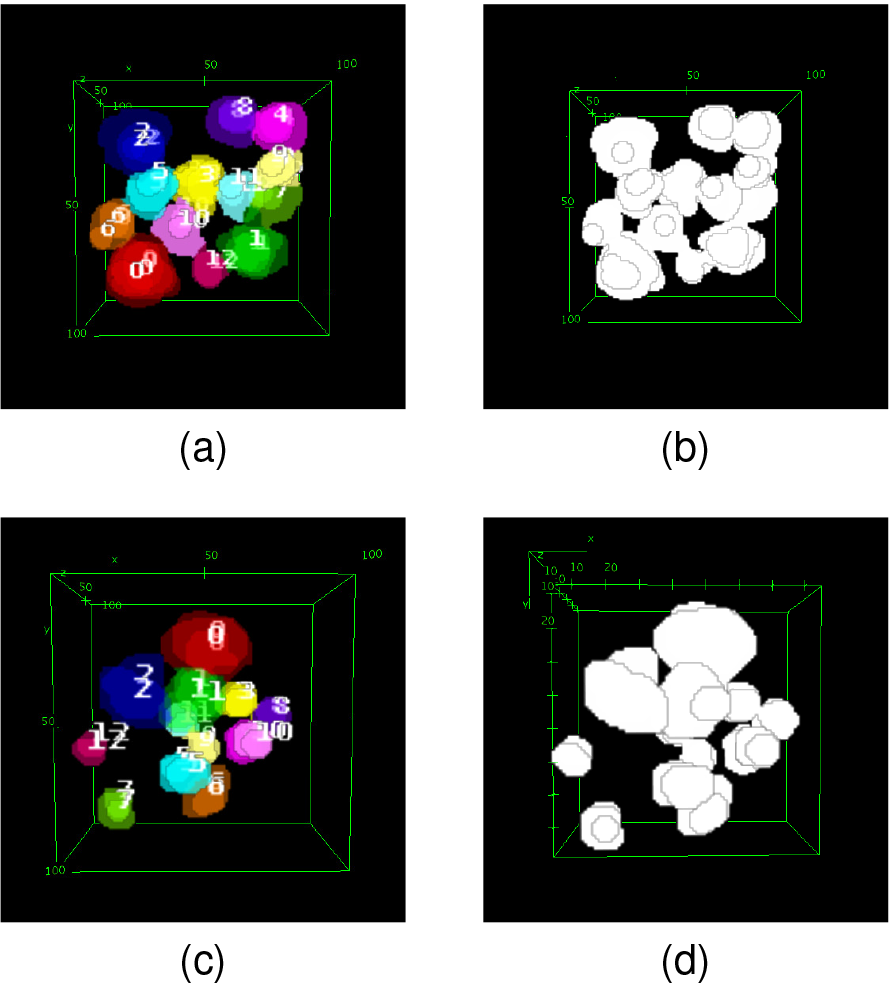
results of the segmentation, 3a,3c, of manually drawn spots 3b,3d, in a top view 3D rendering. Stacks were composed of 6 squared images with a side length of 150 pixels spaced a constant distance of 8.5 pixel equivalents. Each of these images were drawn individually trying to assign high projection overlays to pixel aggregates of contiguous confocal sections. Annotation and color are the same for plane intersections that have been assigned to the same object.

**Figure 4:**
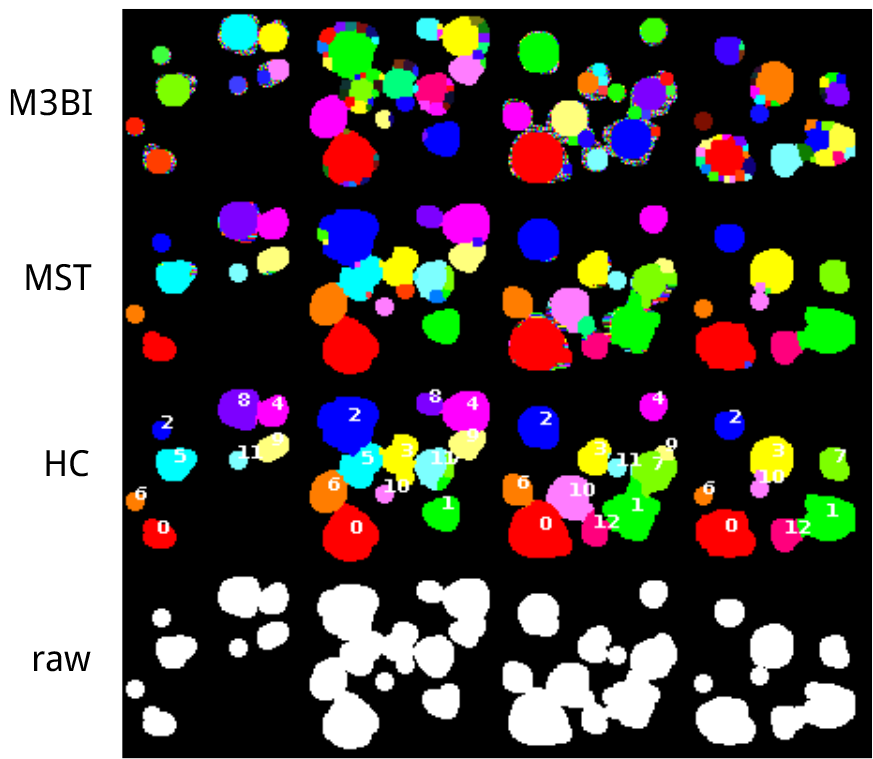
detailed view of the regional merging that resulted in 3a at different stages of the processing. From the top row to bottom, the execution was stopped at a specific stage, and from left to right, each frame corresponds to a different depth within the stack. Only the middle sections from the stack are shown. The two top rows show the result of skipping either all steps after the execution the initial division into maximum 3-balls intersections (M3BI) or after the minimum spanning tree contraction (MST). The third row is the final result after the hierarchical classification (HC) of sets by weight. Bottom row (raw) shows the spot drawing for comparison

**Figure 5:**
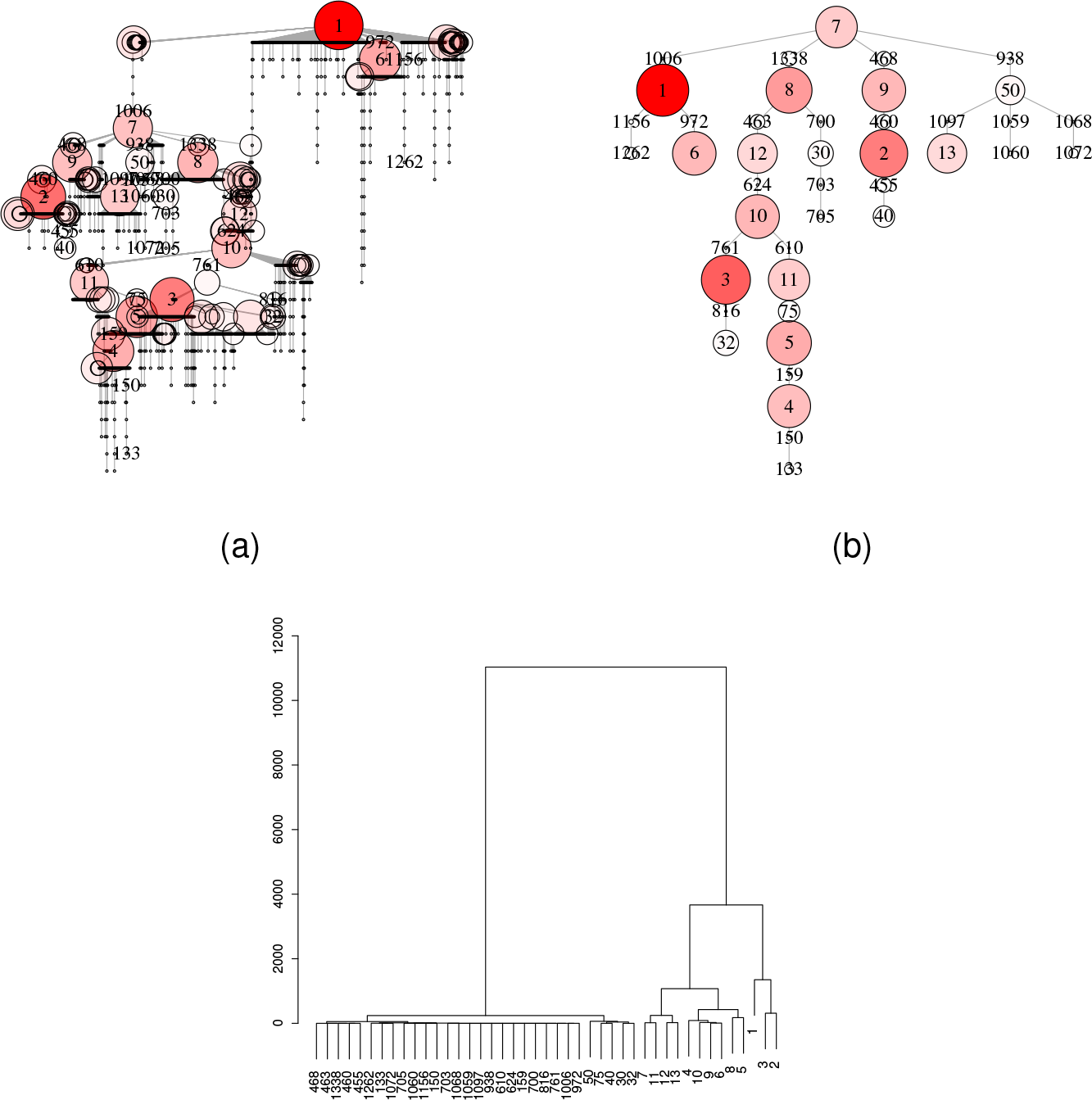
underlying output from the test at 3a at different stages of the execution. 5a, minimum spanning tree (MST) before any contraction is performed. Color intensity and size of vertexes reflects the value of each vertex weight (i.e. ball size). 5b, limit minor that results after the MST contraction. Edges are weighted according to the proposed map 3, 5c, hierarchical classification following the Ward’s minimum variance method. All sets in the right branch almost resemble the final segmentation (i.e. *main sets*). In the final step, voxels from the left branch (i.e. *residual sets*) are assigned to the sets from the right branch according to proximity

We further tested the performance on binarized DAPI stainnings from *Drosophila melanogaster* eye-antenna imaginal discs. *Drosophila* eye discs typically consist on a double layered sac in which one layer is formed by a squamous monostratified epithelium with widely scattered nuclei, i.e. the *peripodial epithelium*. The other layer is thicker and cosists on a pseudostratified epithelium with highly crowded *nuclei*, the so-called *main epithelium*. Cells from this layer are known to undergo differentiation into the mature photoreceptors of *Drosophila* compound eye upon signalling from a moving front wave known as the morphogenetic furrow (MF) [22]. The high overlap of DAPI signal between *nuclei* at the main epithelium makes the task of *nuclei* segmentation challenging for conventional segmentation procedures at this location, e.g. watershed. We wanted to determine wether our segmentation method was able to retrieve the right shape of *nuclei* based on *a)* a rather irregular *nuclei* signal (i.e. DAPI), *b)* a low z-step resolution and *c)* absence of common pre-filtering procedures (e.g. differential of gaussians, DoG). DAPI stainnings were performed in a hypotonic buffer (0.75x PBS), which we notice enhanced nuclear resolution by diminishing signal crowding. The images for the stack were acquired with a Leica SPE confocal laser scanning microscope at a relative resolution of 8.487 equivalent pixels in the z axis and a 405 nanometer excitation laser, which produced 2 or 3 sections per *nucleus* on average. The images were later binarized individually by the Otsu method and processed using iFLIC. In some cases, images were also locally equalized by a CLAHE filter provided by ImageJ before binarization.

The results of this test indicated a quite exact correspondence between each individual *nuclei* and the segmentation (figure 6). One interesting observation is that the segmentation was *noise-robust*, meaning that each white voxel was assigned correctly to one object regardless of their potential elimination by a filter such as DoG. Even when some region was tangential, and so being composed of groups of black voxels embedded in white voxels, the whole group was correctly assigned (figure 6b). In some cases, these tangential regions appeared as misassigned in individual sections. However a three dimensional inspection reveals that in fact these are contact zones of larger segments located up or below the plane, making their assignation ambiguous for the given resolution (figure 6b, white arrows). We expected this ambiguous zones to be reduced at higher z-resolution (i.e. lower z-step). A comparison with the results of a watershed reveals an extensive oversegmentation of the same image without filtering (figure 6b, red arrows).

**Figure 6:**
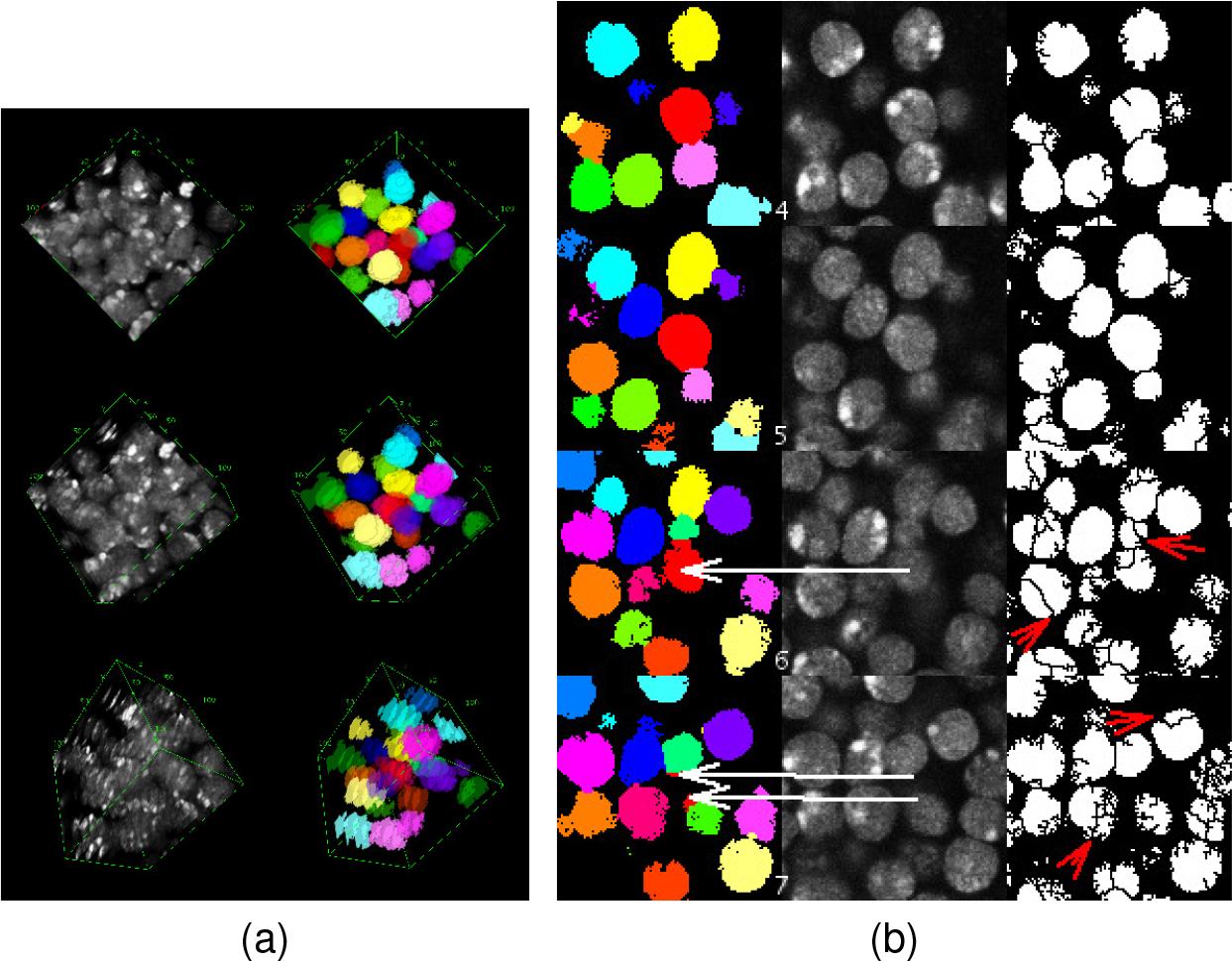
segmentation of a block of cells stainned with DAPI from the ventral flap of a *Drosophila* eye imaginal disc. Sections were acquired with a plane resolution of 179.21 nm·px^−1^ and a z resolution of 1521.0 nm·px^−1^. Pinhole was set so that confocal width covered 1000 nm. approximately. 6a, 3D rendering of the result, identifying the objects in random colouring. 6b Middle sections of the segmentation result. i, iFLIC segmentation; ii, original DAPI signal; iii, results of applying watershed on a binary image without filtering. White arrows point to nuclear segments in contiguous confocal planes that appear to be misassigned; we interpret these regions as ambiguous regions of contact between to objects whose main location is displaced in the z direction. Red arrows point to several oversegmentation examples produced by the watershed

We also wanted to check the effect of *crowding* and *signal overlap* in more controled conditions. Real samples do not offer an objective definition of the shapes and dimensions of the objects (i.e. they usually depend on the judgement of manual segmentation) and critical features such as extension, ellipsoid eccentricity or irregularity of the boundaries cannot be controled individually. Therefore we consider the generation of synthetic samples of objects in three dimensions with different levels of crowding, signal overlap and z-resolutions. Since synthetic samples are generated in a controlled fashion, their shapes, boundaries and dimensions are known *a priori* with no need of manual segmentation. We designed these samples based on intersections of randomly located solid spheres with a radius of 10 voxels and z-steps equal to either 1.25, 2.50 or 5.00 voxels. The total volume of the samples was the same at the three z-steps. Crowding was set by controling object density in the interval of 5 to 80 elements per megavoxel^3^. On the other hand, signal overlap was controled by generating the spheres in two steps. First we produced spheres of radius 10 − *x* at a random location and created an internal representation of this volume. This volume was forbbiden upon selection of a random location for a new object in the next round. The process is repeated until the right amount of objects (according to the selected density) is reached. Second, the spheres at their current location experimented an expansion of *x* pixels beyond their boundaries. The expansion has a random component, since only 1 voxel out of the total allowed neighbour positions (with a upper bound of 26) for each boundary voxel was picked, creating fuzzy borders (figure 8a). The quantity 10 − *x* specifies the minimum potential distance between two sphere centers, hence *x* is a parameter that measures how an expanded object penetrates or *collide* into other object at most, i.e. its *collision index*. A higher density entails a higher probability of collisions since there is less space to locate an equal number of objects; if a higher collision index was set, more objects would collide at higher densities, making it more difficult to identify the spheres. Together with the z-step parameter, an increase in the crowding and the collision index have the general effect of reducing the amount of available information for a correct segmentation, a circumstance that can be used for testing how a particular segmentation method compares to another one. For this assay, a collision index rank of 1 to 6 voxels was used for the comparison of iFLIC with the 3D-watershed implementation provided by the image analysis suit Vaa3D, which includes a prefiltering step [12]. All parameters at the filtering step were reduced in order to minimize the impact on the shape and dimensions of the objects. The segmentation was configured to an adaptive shape-driven watershed which proved to give best results on our synthetic samples. After segmentation, we also trimmed 25 pixels from the border of the images to prevent possible effects that may spoil the results, excluding potential misassigned objects. In order to evaluate the segmentation accurancy, we calculated the mean squared error (MSE) of the minimum shift for the segmented object centroids relative to the true object centroids, i.e. the mean squared difference of distances between each segmented object centroid and the closest object centroid from the synthetic sample. We also determined the MSE of the minimum size shift between synthetic and segmented objects.

iFLIC segmentation came up with a smaller MSE than Vaa3D in all tested conditions except for one case (figures 8b and 8c). MSE for Vaa3D was ~6.45 times higher than iFLIC on average for the centroid shift and ~4.36 times higher on average for the size shift of the closest objects. The higher rate error of the centroid estimation can be attributed to the frequent oversegmentation of watersheds, since the fragmentation of one object leads to several objects bearing a displaced centroid. On the other hand, the higher error rate for size estimation is attributable also to prefiltering, which reduces the amount of voxels around the sets (figure 7). This fact suggests that an improvement of watershed performance by including a prefiltering step occurs at expense of increasing the error rate of the shape and size estimation, therefore reducing the objective value of the watershed as a segmentation method. The two methods also presented a different rate of MSE increase upon reduction of information content of the sample, either by incresing the z-step (i.e. lowering the z-resolution) or collision index (figure 9, table 1). The slope for the regression line was higher when Vaa3D was used in all cases.

**Figure 7:**
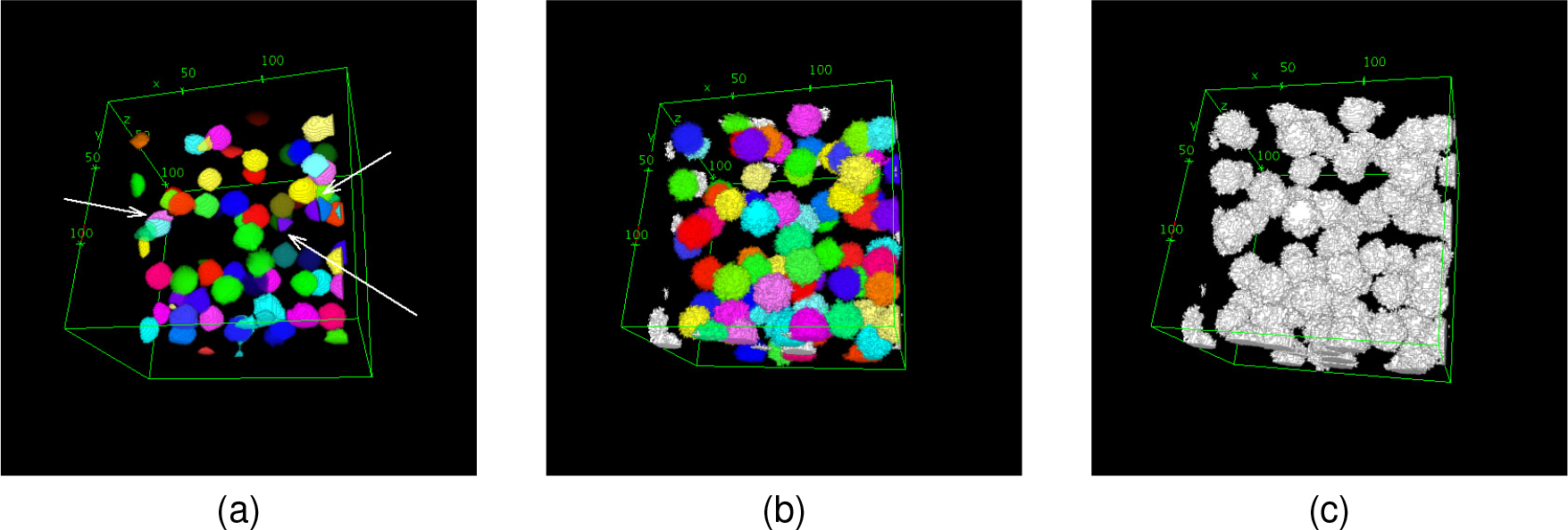
appearance of the segmentation of synthetic samples. 7a, segmentation by Vaa3D suit; 7b, segmentation performed by iFLIC; 7c, original signal. White arrows point to oversegmented instances

**Figure 8:**
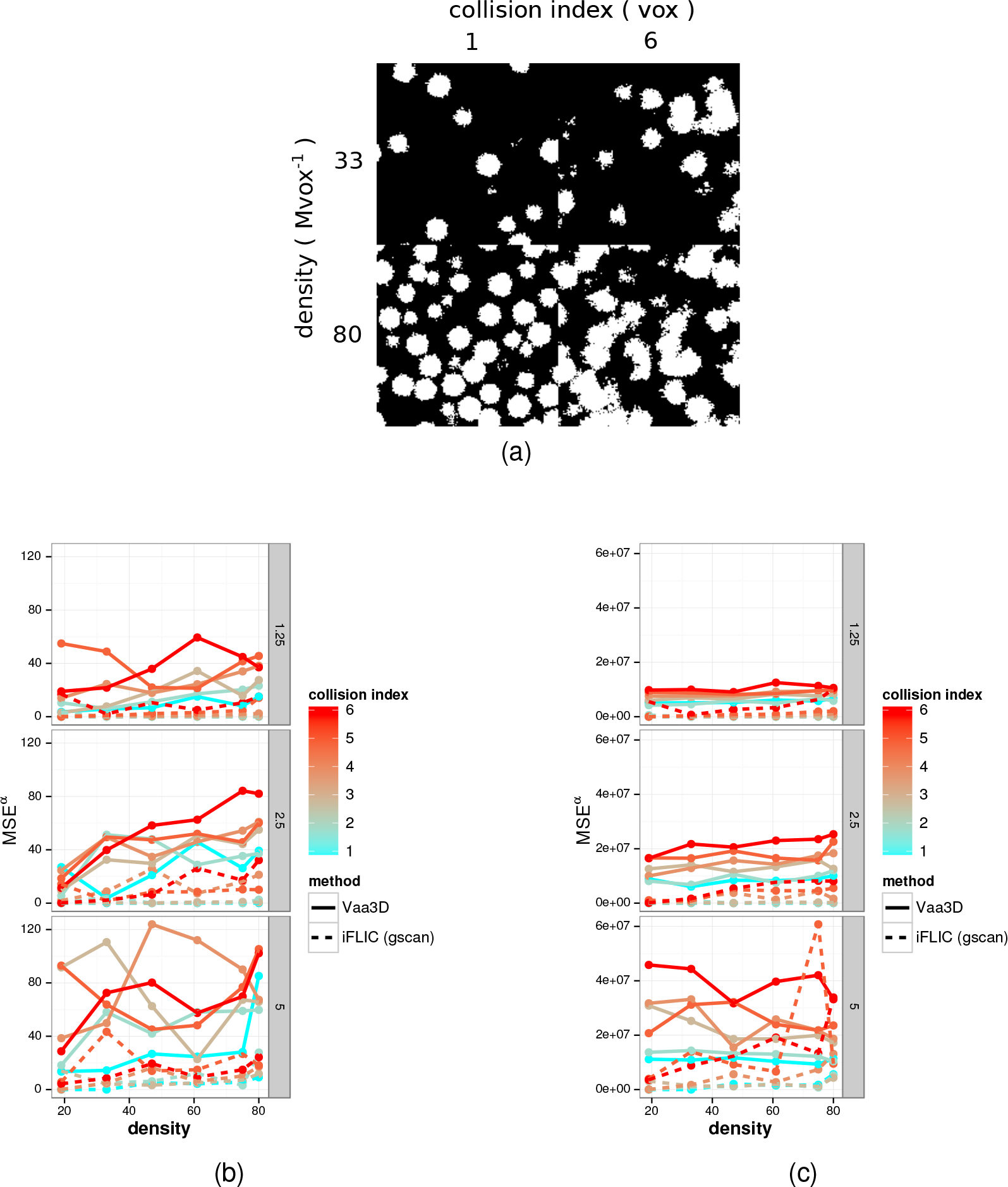
mean squared error (MSE) for the centroid shift and the size shift in synthetic samples for iFLIC and Vaa3D. 8a, four illustrative sections of synthetic samples at different densities and collision indexes. 8b, MSE for the centroid shift. MSE was computed as the distance between the centroids of the closest sets. 8c, MSE for the size shift, i.e. MSE for the size difference of the the closest sets. Panes from top to bottom represent z-steps of 1.25, 2.50 and 5.00 voxels respectively.

**Figure 9:**
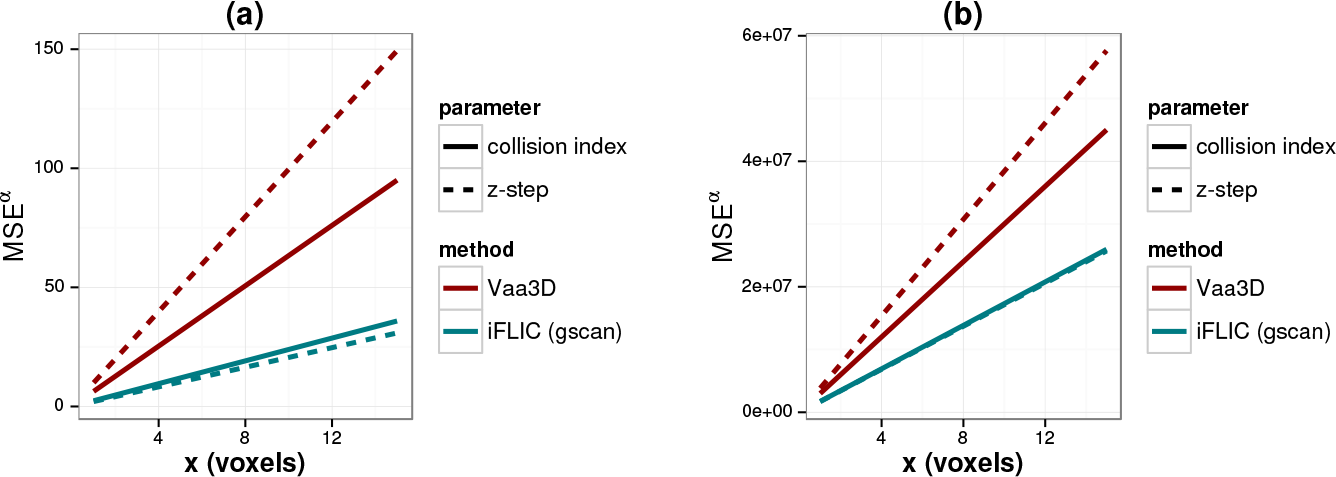
rate of MSE increase for either the centroid shift (a) or the size shift (b) upon increase of z-step or the collision index. Slopes for these lines were extracted from linear regression of the data shown in figure 8.

**Table 1:** mean squared errors for both centroid shift and size shift between synthetic object and segmented objects

| parameter | method | value | MSE (centroid) | MSE (size) |
| --- | --- | --- | --- | --- |
| z-step [vox] | iFLIC (gscan) | 1.25 | 2.67 | 1057417.58 |
|  |  | 2.50 | 6.37 | 1879067.71 |
|  |  | 5.00 | 10.61 | 7222621.74 |
|  | Vaa3D | 1.25 | 23.75 | 7732303.74 |
|  |  | 2.50 | 40.93 | 14235623.66 |
|  |  | 5.00 | 62.05 | 22481245.28 |
| coll.index [vox] | iFLIC (gscan) | 1.00 | 1.36 | 613504.78 |
|  |  | 2.00 | 3.60 | 653948.59 |
|  |  | 3.00 | 2.71 | 795361.71 |
|  |  | 4.00 | 7.95 | 2558537.68 |
|  |  | 5.00 | 11.17 | 7306528.11 |
|  |  | 6.00 | 12.51 | 8390333.19 |
|  | Vaa3D | 1.00 | 22.80 | 8306457.88 |
|  |  | 2.00 | 32.78 | 9122878.09 |
|  |  | 3.00 | 41.65 | 14121086.46 |
|  |  | 4.00 | 50.26 | 15934585.06 |
|  |  | 5.00 | 52.22 | 17474589.95 |
|  |  | 6.00 | 53.75 | 23938747.91 |
| density [Mgvox<sup>-1</sup>] | iFLIC (gscan) | 19.00 | 3.37 | 1020595.29 |
|  |  | 33.00 | 4.89 | 1838073.04 |
|  |  | 47.00 | 6.59 | 2748361.68 |
|  |  | 61.00 | 5.76 | 2920773.84 |
|  |  | 75.00 | 7.18 | 6212719.42 |
|  |  | 80.00 | 11.50 | 5577690.80 |
|  | Vaa3D | 19.00 | 26.96 | 14915056.99 |
|  |  | 33.00 | 39.47 | 15571806.97 |
|  |  | 47.00 | 40.69 | 13983523.83 |
|  |  | 61.00 | 43.38 | 14674116.82 |
|  |  | 75.00 | 47.00 | 14954426.64 |
|  |  | 80.00 | 55.95 | 14799414.11 |

The estimation of the total amount of sets was also affected by an increase of critical parameters (figure 10). Despite that both Vaa3D and iFLIC performed equally well at a high z-resolution (i.e. 1.25 voxels), when the z-step was increased Vaa3D started to deviate from the right estimation dramatically. A more detailed observation reveals that iFLIC’s loss of precision was mainly observed as both collision index and density increased, introducing a bias towards undersegmentation, i.e. an underestimation of the amount of objects. Vaa3D presented a deep bias in both directions depending on the collision index or density, but mainly on the z-step as mentioned. An undersegmentation bias was observed when the z-step was high, which turned to a slight oversegmentation as z-step decreases. This results point to a much lower impact of the z-resolution when iFLIC is used.

**Figure 10:**
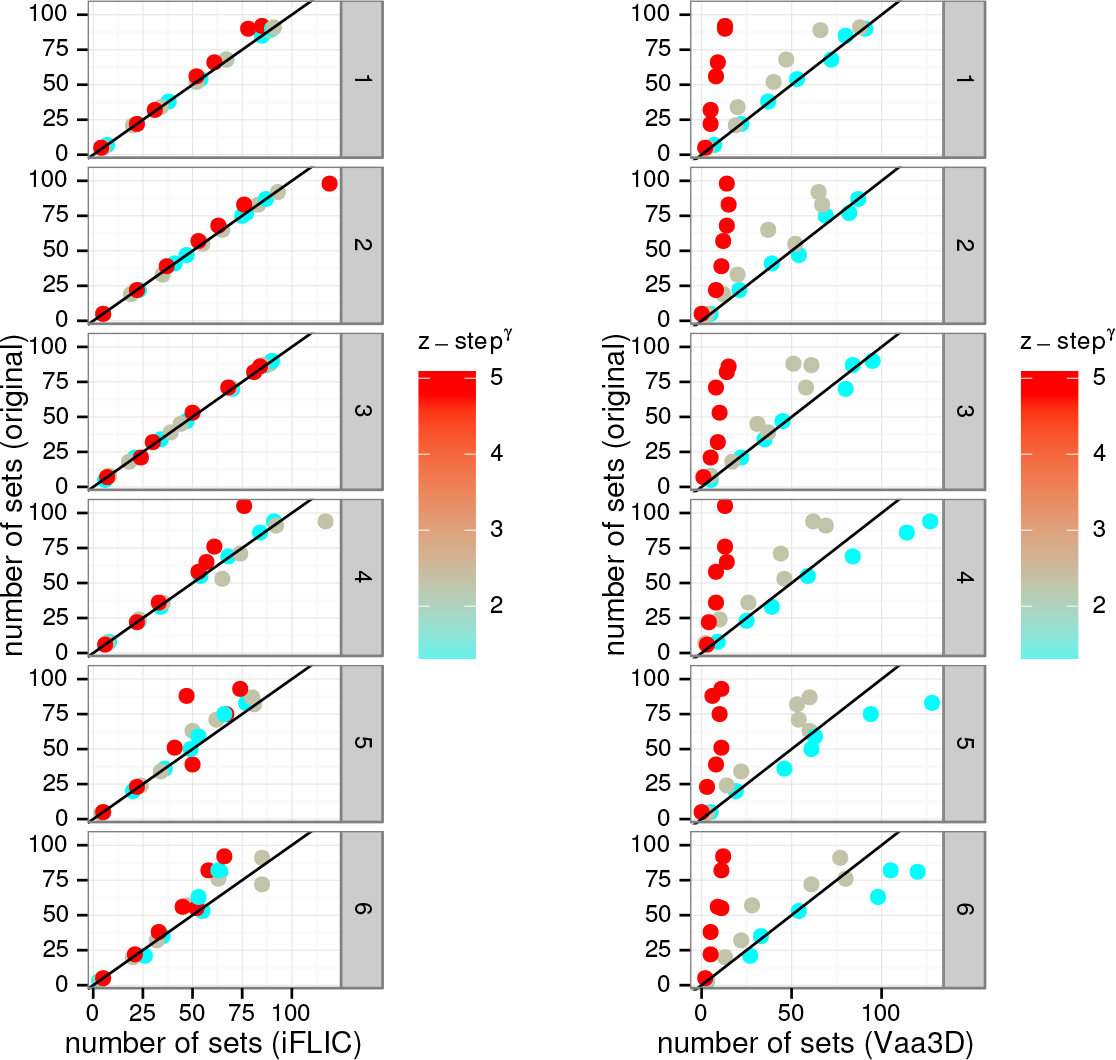
prediction of the right number of sets using iFLIC and Vaa3D methods. Predicted sets are shown in the *x* axis; *y* axis shows the true number of sets for which the prediction was performed, hence better predictions tend to get close to the main diagonal (black line), undersegmentation would tend above the main diagonal, and below if the sample is oversegmented. Each frame, starting at the top, represent increases of 1 voxel in the collision index. Left pane, iFLIC; right pane, Vaa3D.

## 4 discussion

In this work we have proposed a new method for segmentation based on topological concepts, graph theory definitions and hierarchical clustering. We also developed a software (iFLIC) based on these ideas with the aim of testing the adequacy in real and synthetic samples displaying typical features from confocal laser scanning microscopy (LCSM), such as noisy and irregular signal and a relatively low resolution in the z direction. We have proved that the proposed method copes better with these conditions than watershed based methods in 3D. We also determined that for an acceptable performance (i.e. a significant reduction of oversegmentation), watershed methods require the use of prefiltering steps that are commonly found in available software such as Vaa3D, which in turn spoil the retrieval of the objects that these methods aim at for segmenting, e.g. introducing variations on the size, centroid and amount of predicted objects. Our method has rendered better results with no use of filtering before the execution. Since filtering, such as median or gaussian, are usually affected by some user-set parameters (e.g. kernel size) it also introduces a source of subjective variability that our method does not, making it reproducible between individuals.

Another important difference between watersheds and our method is the fact that the former are designed to reveal the boundaries between objects that share some degree of overlap. In flat images this usually leads to the removal of boundary pixels such that this outlining would facilitate segmentation by flooding.

However we think this is only a valid approach for flat images whose object size is large enough relative to its resolution and the contribution of the boundary pixels to the total proportion of pixels from the object is low. This situation changes when image stacks coming from LSCM are used because the resolution in the z direction is usually lower. Following the same procedure, a removal of boundary voxels, whose contribution is much higher regarding the total size of the objects, impacts more dramatically the estimation of important object features, as we proved by the high change in performance of the prediction of the number of objects (figure 10). We interpret this effect as a way of focus: segmentation problems are better set up in terms of pixel (voxel) assignation to objects or pixel (voxel) classification, instead of frontier pixels (voxels) detection and removal. We followed this rationale by applying a regional merging method for which the final segmentation arises naturally from the definition of *limit minor*, showing that small unclassifiable regions tend to correspond to those boundaries that watersheds are designed to detect. We showed that a hierarchical classification can extract this unclassified regions (i.e. residual sets), being integrated into the main sets and leading to a satisfactory resolution in terms of segmentation with some unexpected and interesting properties, e.g. noise-robustness. However we did not assumed that Fano’s factor could be the best cost function for this classification. On the contrary we tested cost functions for common hierarchical classification procedures, such as SLINK [23] and Ward’s minimum variance methods [19] (figure 5c) with a quite well approximation to the result we expected. Nevertheless when Fano’s factor was used, the result was exact in a much wider set of conditions. Since Fano’s factor is considered a measure of the noise-to-signal ratio, this observation is congruent with the interpretation that residual sets are actually some sort of “noise”, justifying that the voxels from the residual sets were delivered to the main sets by proximity.

We also stress the importance of computing the MST from the initial graph. After testing the effect of finding a limit minor from *G* without computing a MST, we noticed that the result tended to be undersegmented. We interpreted this fact as a need for restricting the set of edges that composes *G*, i.e. finding a subgraph of *G* that favoured the intended segmentation. Of course this does not mean that the best subgraph has to be a spanning tree; however finding a spanning tree seems convenient from a practical point of view since fast standarized methods for its computation already exists (e.g. Prim’s, Kruskal’s) [21, 24]. We do not demonstrate mathematically the adecquancy of our edge weighting function (i.e. equation 3), but we proved its usefulness after the results obtained with its application.

One drawback is that, unlike watersheds, which are optimized to run in a *O* (*n*) time complexity, our algorithm present a lower bound of *O* (min {5〈*b*〉, *n* − 1}*n*) time complexity at the step of graph building. All remaining steps are optimized beyond this point since Prim’s and DFS usually handle a low number of sets relative to the size of the images and also run in parallel for each component of the initial graph. However we think our current algorithm could be optimized at graph building by dividing into balls individual components found by flood fill, which would also allow a parallelization from an earlier step.

Finally we point out that methods such as ours offer several practical advantages, since if it performs well at low z-resolution, image acquisition times are reduced, reducing concomitantly photobleaching of the samples and also the data size. Moreover, increasing the resolution naturally results in a increase of the computational time whatever method is used. These facts have urged us to exploit as much as possible methods with a higher potential for information management, for which we hope to have contributed with the ideas presented in this work.

## Footnotes

1 More exactly this means that *G* is labeled at its vertexes according to some value associated with each ball (e.g. *w*)

2 in very rare cases a ball is composed by just one voxel with 6 potential neighbours

3 this measure is performed in an equal volume but, since not all samples bear the same z-step, different voxel dimension in the z direction is found. For this reason when we talk about voxel we refer to cubes with an equivalent pixel length in the z direction, as in this case. This ensure that densities are consistently equal after z-step variation

## A Java code for *limit minor* search

Here we present the complete Java implementation for *limit minor* search included in the gscan plugin, which is based on the algorithm 1.
the two major classes are:

1. `ZConnectedSet`. this class stores the voxels of individual sets of points. It also includes a separated boundary voxel list and total size (through the method `getSpaceExtension`) as parameters, and boundary voxel and centroid calculation methods.
2. `GZNSet`. this class extends `ZConnectedSet` capabilities for graph definition. It includes a parameter (i.e. S) representing a `ZConnectedSet` instance associated to a graph vertex and a `Set` (backed up by a hash table) of `GZNSet` instances that act as first order nearest neighbours (1-NN) vertexes. Some other static methods are found in this class, e.g. contact surface calculation.

These two classes operate within a `GClusterProcessor` class, which extends the Thread class and overrides its `run` method. This allows to implement the limit minor search algorithm in instances that may run in parallel. Individual instances of the `GClusterProcessor` class are all operated within the class `GraphScanner`, whose constructor accepts user parameters from the `iCommand` interface, the common interface of all iFLIC plugins.

Other graph definitions are required for the implementation of DFS and Prim’s for which specific graph conversor methods are also provided within the `GraphScanner` class.

The *limit minor* code is implemented within the `GClusterProcessor` class (method `ContractGraph`, code 1). The header defines a prefixed set of memory allocations, one queue, `PQ`; and two stacks, `T` and `N`, and accept a regular list of `GZNSet` objects, i.e. `zsS. PQ` stores the priority queue of the algorithm 1; the other two stacks are temporal allocations for nonempty sets of `zsS`, i.e. `T`, and the 1-NN of the current `GZNSet` instance, i.e. `N`. It calls to two external static methods, one for testing whether an edge is degree reducing (i.e. `getDegreeReduction`, code 2) and other for the combination of two sets of points upon contraction, (i.e. `combineGZNSets`, code 3).

The flag `b` controls if some contraction has occured in the previous cycle and hence, let the program “check” one more time whether some other contraction is possible.

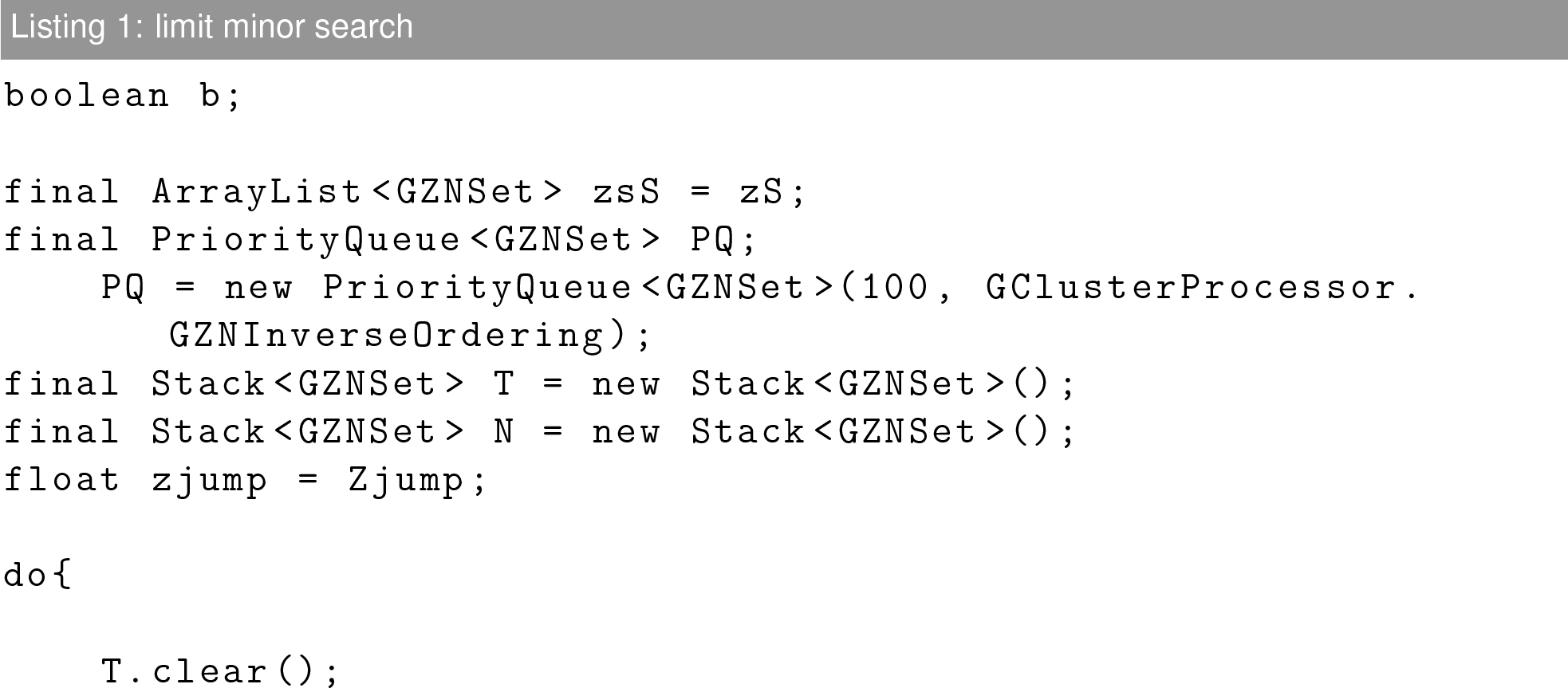

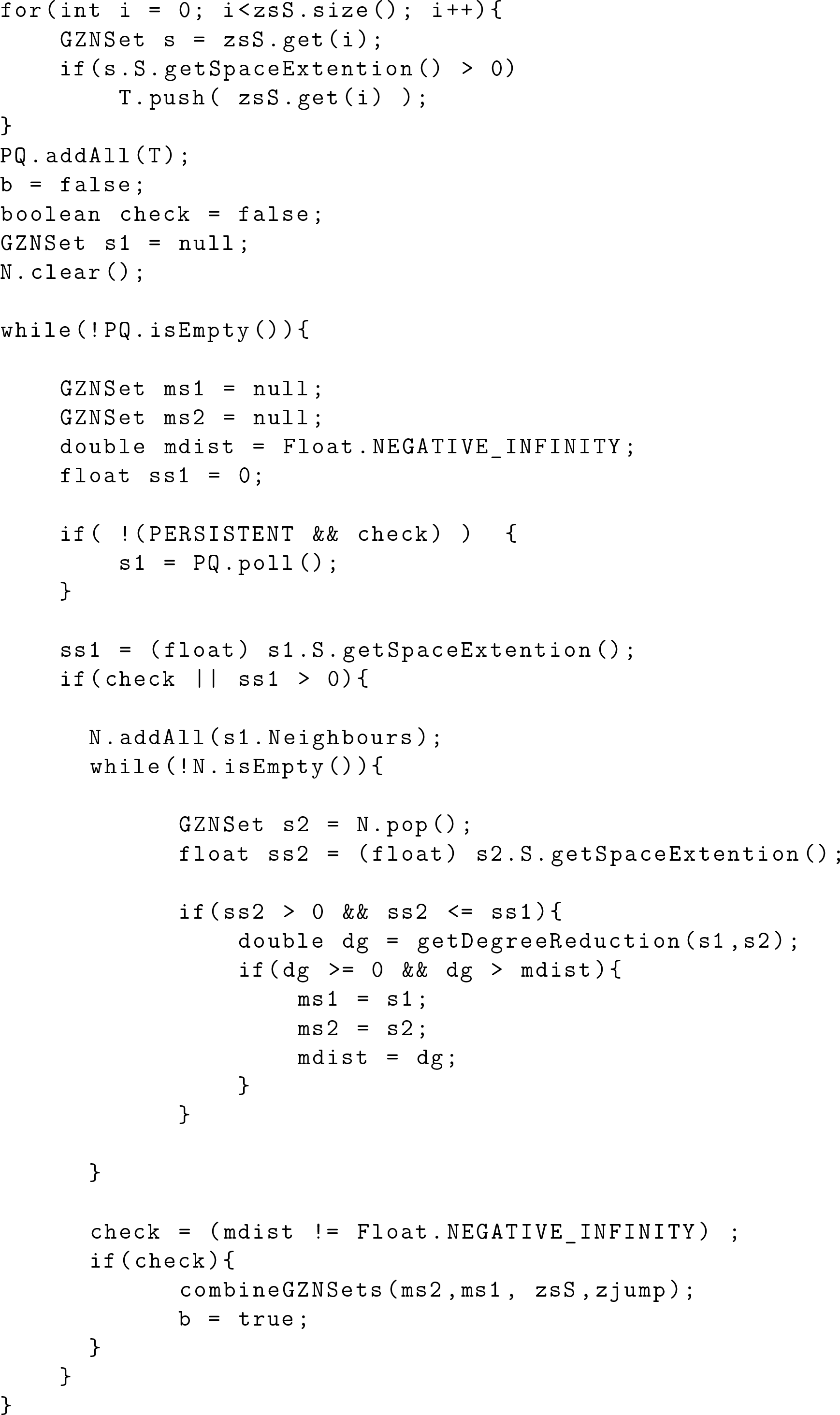

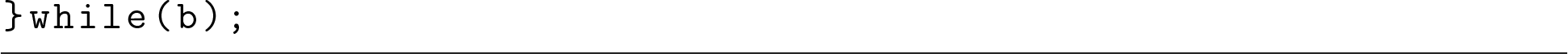

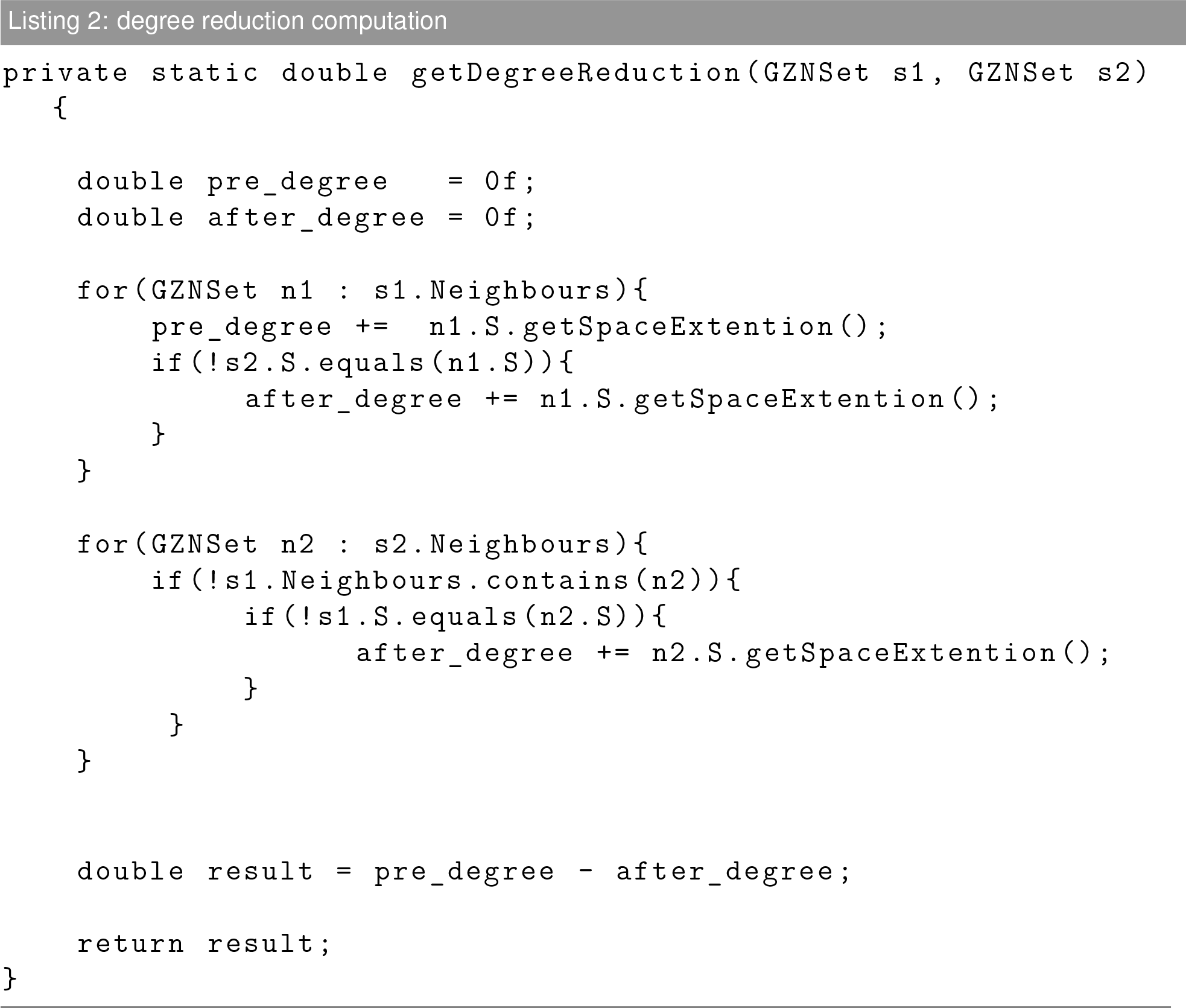

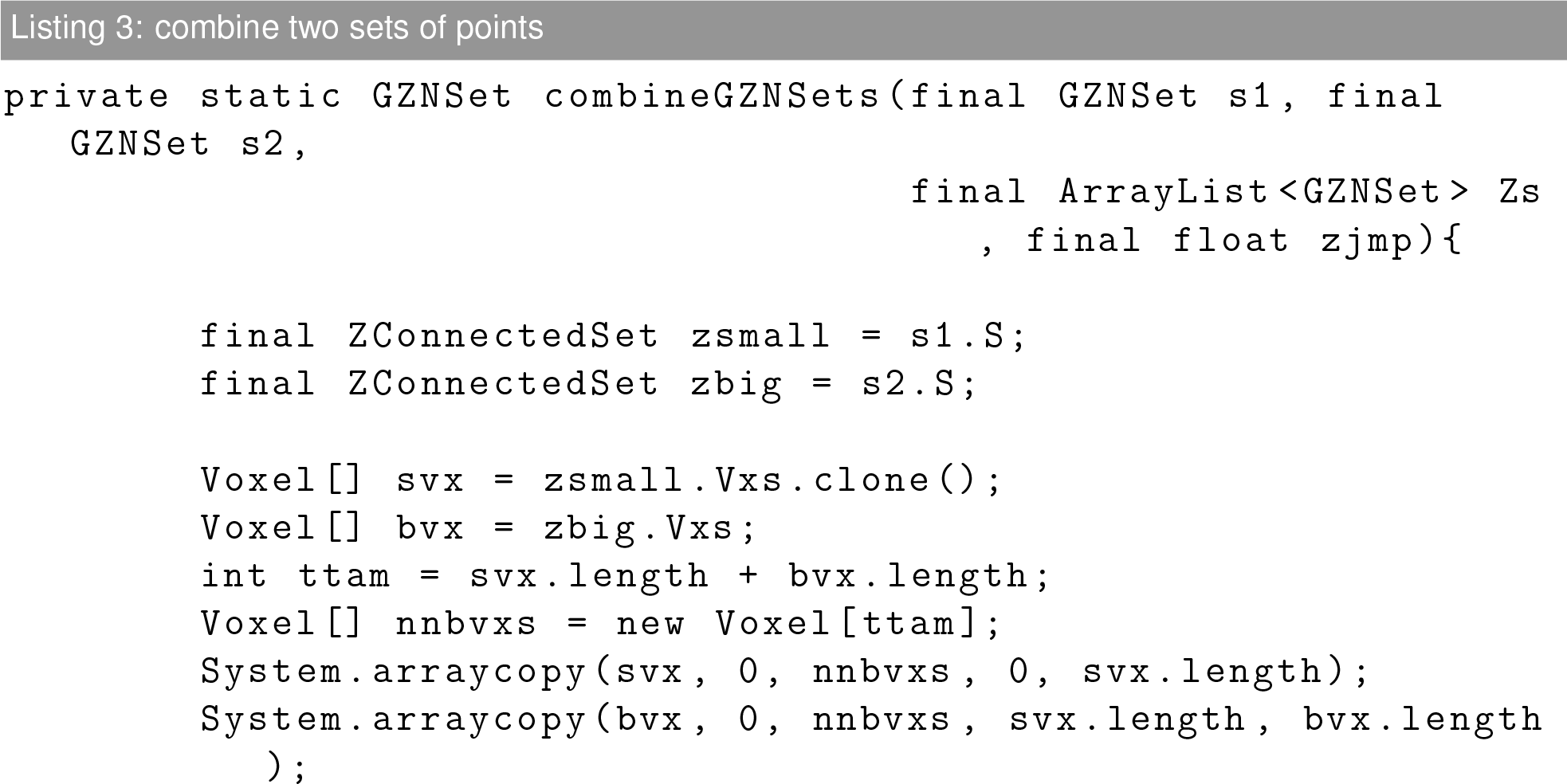

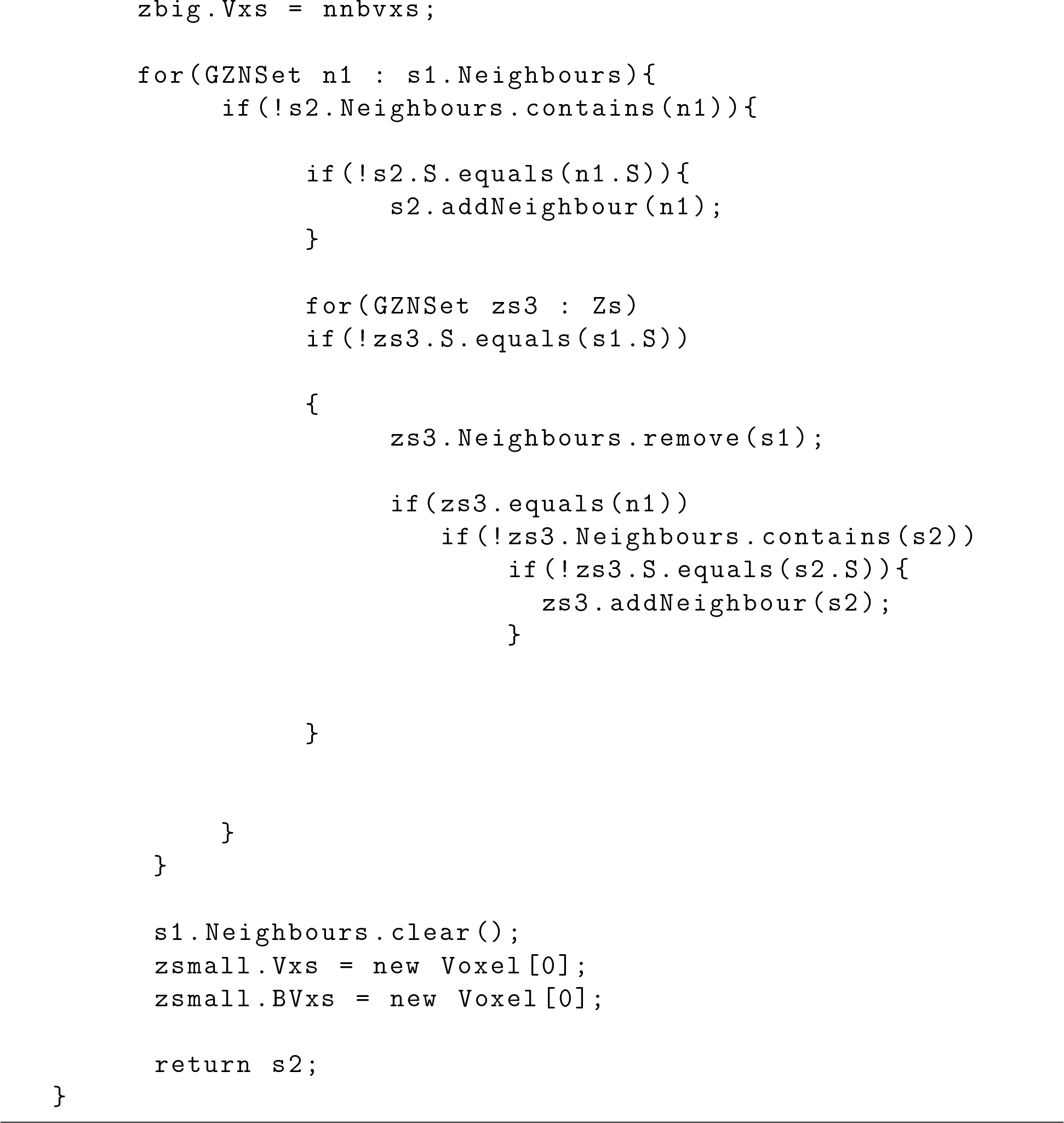

